# Boron deficiency-induced root growth inhibition is mediated by brassinosteroid signalling regulation in *Arabidopsis*

**DOI:** 10.1101/2019.12.12.874222

**Authors:** Cheng Zhang, Mingliang He, Sheliang Wang, Liuyang Chu, Chuang Wang, Ningmei Yang, Guangda Ding, Hongmei Cai, Lei Shi, Fangsen Xu

## Abstract

Brassinosteroid (BR) is a pivotal phytohormone involved in regulating root development. Boron (B) is an essential micronutrient for plant growth and development, and root growth of plants is rapidly inhibited under B deficiency condition, but the mechanisms are still elusive. Here, we demonstrate that BR plays crucial roles in these processes. We identify BR-related processes underlying B deficiency at the physiological, genetic, molecular/cell biological and transcriptome levels, and provide strong evidences that B deficiency can affect BR signalling, thereby altering root growth. RNA-sequencing analysis reveals a high co-regulation between BR-regulated genes and B deficiency-responsive genes. We found that low B negatively regulates BR signalling to control BR signalling-dependent root elongation, *bes1-D* exhibits insensitivity to low B stress, and *bri1-301* mutants fails to respond to B depletion. Exogenous eBL application can rescue the inhibition of root growth under B deficiency condition, and application of BR biosynthesis inhibitor BRZ aggravates root growth inhibition of wild-type under B deficiency condition. B deficiency reduces the nuclear signal of BES1. We further found that B deficiency reduces the accumulation of brassinolide (BL) by reducing *BR6ox1* and *BR6ox2* mRNA level to down-regulate BR signalling and modulate root elongation. Altogether, our results uncover a role of BR signalling in root elongation under B deficiency.

**One-sentence summary:** B deficiency reduces the accumulation of brassinolide by reducing *BR6ox1* and *BR6ox2* mRNA level to down-regulate BR signalling and modulate root elongation.

## INTRODUCTION

Warington gave the first definitive proof of boron (B) requirement for a higher plant in 1923 (Warington, 1923). Since then, B has been identified an essential micronutrient for higher plants. The main known function of B in higher plants is to cross-link the apiosyl residues of two neighbouring monomeric RG-II (monorhamnogalacturonan-II, mRG-II) molecules for forming a dimeric B-dRG-II pectin complex (Kobayashi *et al*., 1996). This role of B in cell walls was confirmed by the *Arabidopsis mur1* dwarf mutant with brittle stems, which could be rescued with added B (O’Neill *et al*., 2001).

Boron deficiency is a prevalent micronutrient deficiency in agricultural production worldwide, which leads to crop yield loss and quality decline (Shorrocks, 1997). B deficiency leads to a wide range of morphological, and physiological symptoms, such as apical growth inhibition, leaf expansion reduction, terminal bud necrosis, flower abortion and fruit shedding (Goldbach, 1997; Brown *et al*., 2002). Among these symptoms, one of the earliest responses to B deficiency for plants is the inhibition or cessation of root elongation (Dell *et al*., 1997). Reduction of root growth under B deficiency has been reported in many dicots, including *Arabidopsis, Brassica napus* and orange (Takano *et al*., 2006; Hua *et al*., 2016; Wu *et al*., 2017). Graminaceous plants demand less B, but gramiaceous mutants of B transporter genes display conspicuous inhibition of root elongation (Nakagawa *et al*., 2007; Durbak *et al*., 2014; Shao *et al*., 2018). Two protein families are involved in B transport, the NIP influx channels and the BOR efflux transporters (Miwa & Fujiwara, 2010). However, the mechanisms through which B deficiency inhibits root growth are still enigmatic.

Brassinosteroids (BRs) are a class of polyhydroxylated steroid phytohormones in plants. BRs regulate a wide range of physiological and developmental processes including cell elongation, cell division, senescence, root hair initiation, stomata development, vascular differentiation, reproductive development and photomorphogensis (Clouse & Sasse, 1998; Divi & Krishna, 2009; Wei & Li, 2016). BRs also interact with many other hormones and environmental factors to modulate plant growth and development (Nemhauser *et al*., 2006; Zhang *et al*., 2009; Yang *et al*., 2011). In addition, BRs promote the expression of cell wall synthesis genes and affect the expression of P450 genes involved in BR biosynthesis and catabolism (Goda *et al*., 2002). BRs are synthesized from phytosterols through multiple, mostly oxidative reactions. Campesterols (CR) is thought to be the first molecule specifically for the BR biosynthesis, which is firstly converted to campestanol (CN), then to castasterone (CS), and finally to brassinolide (BL) (Fujioka *et al*., 1995; Choi *et al*., 1996; Choi *et al*., 1997). Several cytochrome P450 enzymes play important roles in the BR biosynthesis, such as DEETIOLATED2 (DET2), CONSTITUTIVE PHOTOMORPHOGENESIS AND DWARFISM (CPD), ROTUNDIFOLIA3 (ROT3), CYP90D1, DWARF4 (DWF4), BRASSINOSTEROID-6-OXIDASE1 (BR6ox1), and BR6ox2. CPD catalyzes C-23 hydroxylation of BRs (Ohnishi *et al*., 2012), and DWF4 is contribute to multiple C-22 hydroxylation (Choe *et al*., 1998). Both BR6ox1 and BR6ox2 catalyze multiple C-6 oxidation, while BR6ox2 catalyzes the lactonization of CS to BL (Bishop *et al*., 1999; Kim *et al*., 2005). Some artificial chemicals inhibitors of BR biosynthesis have been reported. For example, brassinazole (BRZ) can directly inhibit the catalytic activity of DWF4 (Asami *et al*., 2000).

BRs are perceived at the plasma membrane by three leucine-rich repeat receptor-like kinase (LRR-RLKs) BRASSINOSTEROID INSENSITIVE 1 (BRI1) (Li & Chory, 1997). BRs bind to the extracellular leucine-rich repeat domain of BRI1, which phosphorylates BRI KINASE INHIBITOR 1 (BKI1) as a downstream negative regulator of BRI (Wang & Chory, 2006), resulting in the release of BKI1 from the intracellular BRI1 kinase domain, allowing BRI1 to recruit its co-receptor BRI1-ASSOCIATED RECEPTOR KINASE 1 (BAK1), another LRR-RLK (Li *et al*., 2002; Nam & Li, 2002). At downstream of BRI1 and BAK1, members of the BRASSINOSTEROID SIGNALING KINASE (BSK) cytoplasmic kinase family initiate a phosphorylation-and-dephosphorylation cascade (Kim *et al*., 2009). Reversible phosphorylation and dephosphorylation cascade leads to nuclear accumulation of two unphosphorylated transcription factors, BRASSINAZOLE-RESISTANT 1 (BZR1) and BRI1-EMS-SUPPRESSOR 1 (BES1) (Wang *et al*., 2002; Yin *et al*., 2002). BZR1 and BES1 can regulate thousands of target genes to mediate plant growth and development (Sun *et al*., 2010; Yu *et al*., 2011). The hypophosphorylated state of *bes1-D* and *bzr1-D* dominant mutants results in constitutive activated BR signalling and constitutive BR responses (Wang *et al*., 2002; Yin *et al*., 2002).

BRs regulate growth and development of plant roots in a dose-dependent manner (Clouse *et al*., 1996; Mussiget *et al*., 2003). Application of low concentration of BL can facilitate root growth, but high concentration of BL inhibited root elongation (Gonzalez-Garcia *et al*., 2011). Mutants impaired in BR biosynthesis, such as *cpd* and *det2*, showed much shorter roots compared to those of wild-type plants (Chory *et al*., 1991; Fujioka *et al*., 1997). Both *bri1* and *bak1* which were the mutants defective in BR signal transduction, also showed significantly shortened roots (Clouse *et al*., 1996; Li *et al*., 2002). Recent studies found that BZR1 and BES1 activity conferred to root’s insensitivity to Pi deprivation through repressing LPR1 expression and affecting Fe distribution (Singh *et al*., 2014; Singh *et al*., 2018). It was also found that BSK3 as a major determinant for primary root length response to low N condition (Jia *et al*., 2019).

Here, we report that B deficiency impacts on BR-dependent root growth through reducing the accumulation of BL. We show that B deficiency results in reduced root meristem size and mature cell length. The *bes1-D* constitutive BR response mutant showed significantly longer primary root than wild-type under B deficiency condition, while *bri1-301* failed to respond to B depletion. B deficiency down-regulated *BR6ox1* and *BR6ox2* to reduce the accumulation of BL to inhibit BR-dependent root growth. Overall, our data highlight the important role of BR signalling in the adaptation to B deficiency condition.

## RESULTS

### Influence of B deficiency on root growth

To evaluate the impact of B depletion upon root growth in *Arabidopsis*, we performed a root sensitivity assay by monitoring primary root growth and developmental parameters of roots in medium of gradient B concentrations. Primary root length of wild-type plants (En2) was significantly inhibited when B concentration was decreased from 100 μM to 0.05 μM (Fig. 1A and B). The root growth was decelerated rapidly with the decrease of B concentration within 3 days after germination (Fig. 1C).

**Figure 1.**
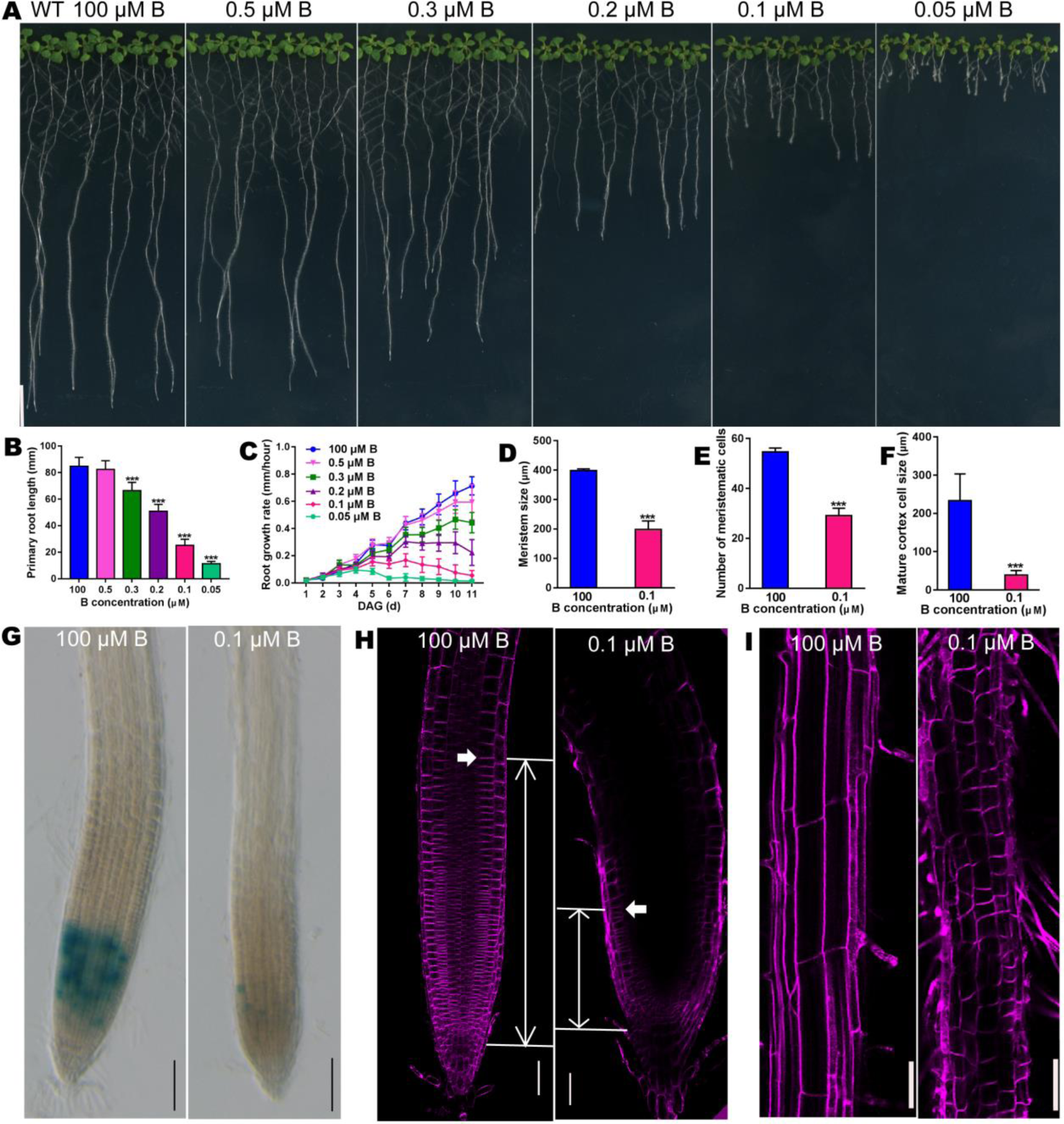
B deficiency impacts on root elongation. **(A)** Phenotype of wild-type plants grown at decreasing concentrations of B for 11 d. Scale bar, 10 mm. **(B)** Primary root lengths of wild-type plants grown at decreasing concentrations of B for 11 d (mean ± s.d., n = 30). The asterisks indicate a statistically significant difference (*t*-test, P < 0.001). **(C)** Root grown rate of wild-type grown at decreasing concentrations of B for 11 d (mean ± s.d., n = 30). DAG, Days after germination. **(D-F)** Quantification of meristem size **(D)** (mean ± s.d., n>6), number of meristematic cells **(E)** (mean ± s.d., n>6) and mature cortex cell size **(F)** (mean ± s.d., n>40) of wild-type root grown as in **(H)** and **(I)**. The asterisks indicate a statistically significant difference (*t*-test, P < 0.001). **(G)** The expression of *pCYCB1;1:GUS* plants grown on MGRL media for 6 d under 100 and 0.1 μM B conditions. Scale bar, 50 μm. **(H)** Confocal microscopy image of 6-day-old wild-type. Arrows represent the meristem boundary where cortical cells double their size. Scale bar, 50 μm. **(I)** Confocal microscopy image of wild-type root grown as in **(A)**. Scale bar, 100 μm.

It is known that the root growth inhibition is the result of a decrease in cell size and/or cell number. To investigate whether the inhibition under B deficiency condition results from the decreased cell number and/or cell size, we measured the number of meristematic cells and the length of fully elongated root cells. Roots of wild-type grown under B deficiency displayed reduced root meristem size, reduced number of meristematic cells (Fig. 1D, E and H), and decreased expression of the *CYCB1;1*, which can be regarded as the maker gene of cyclin cell cycle (Fig. 1G). A significant decrease of mature cell length was also found for plants grown at B deficiency condition (Fig. 1F and I). Taken together, these observations indicate that B deficiency results in reduced number of meristematic cells, root meristem size and mature cell length in roots, and thus reduces root growth.

### Genome-wide root responses to B deficiency

To shed light on the pathways mediating root growth responses to B deficiency, we performed whole-genome RNA sequencing (RNA-seq) analysis to compare root transcriptomes of 9-day-old wild-type plants grown under adequate and depleted B conditions. We found that 4,441 genes showed significantly differential expression in wild-type plants between the depleted B condition versus adequate B condition (Fig. 2B), 2,360 of these genes were strongly (>2-fold) up-regulated or down-regulated (Supplemental Table S1). The top gene ontology (GO) terms (ranked by Correct p-Value) among the differential expression genes showed that B deficiency directly and/or indirectly influences a range of biological processes, including oxidation-reduction, cell wall organization, lipid metabolic and steroid biosynthetic process (Fig. 2A). Increased H_2_O_2_ fluorescence and O_2_^.-^ fluorescence in the root with the fluorescent probes 3’-(*p*-hydroxyphenyl) fluorescein (HPF) and dihydroethidium (DHE) were observed under B deficiency (Supplemental Fig. S1).

**Figure 2.**
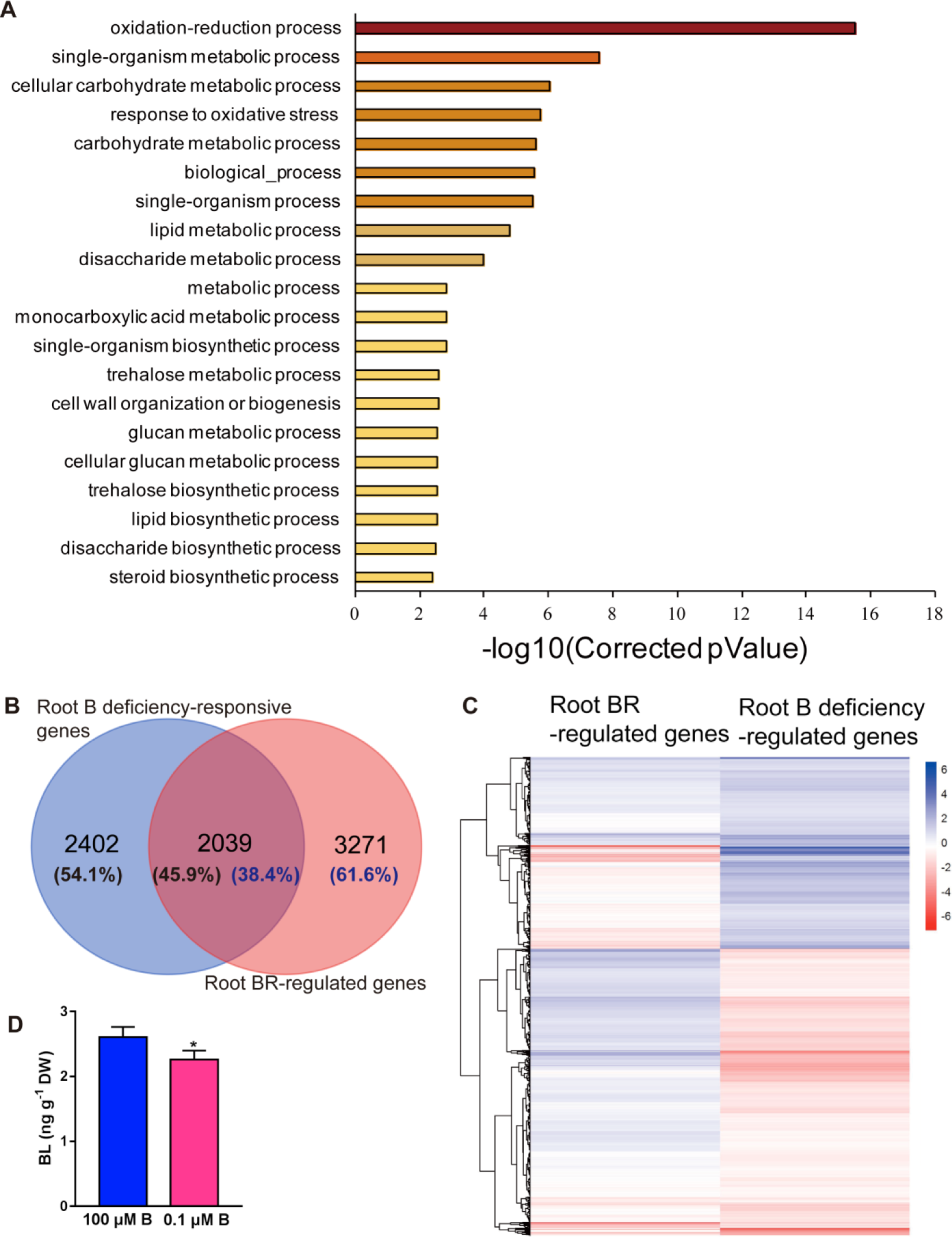
Genome-wide responses to boron deficiency in root. **(A)** Top 20 (ranked by false discovery rate (FDR < 0.05)) significantly enriched GO terms derived from genes that are regulated by B deficiency. **(B)** Venn diagrams representing the overlap between B deficiency-regulated genes and published BR-regulated genes in roots. **(C)** Hierarchical cluster analysis of log2 fold change value of BR-B deficiency co-regulated genes. **(D)** The BL content of wild type En2 under 0.1 and 100 μM B conditions.

Since several recent studies demonstrated that BRs played an important role in root growth (Gonzalez-Garcia *et al*., 2011; Hacham *et al*., 2011; Chaiwanon *et al*., 2015; Jaillais *et al*., 2016), we compared our B deficiency-responsive genes with a recent root-specific BR-regulated gene data set generated by RNA-seq (Chaiwanon *et al*., 2015). Interestingly, 45.9% of the genes regulated by B deficiency were also regulated by BRs in the root (Fig. 2B and Supplemental Table S2), indicating a high degree of co-regulation between them. Furthermore, we found that more than 60% of the co-regulated genes responded to B deficiency and BRs in opposite directions (Fig. 2C and Supplemental Fig. S2A). It has been reported that activated BR signalling appears to inhibit at least five BR biosynthetic genes, as well as *BRI1* through negative feedback, while positively regulated downstream BR signalling by activating transcription of *BSU1* and inhibiting *BIN2* (Tanaka *et al*., 2005; Sun *et al*., 2010). Indeed, *BRI1, BAK1, BIN2* and *BZR1* were accumulated more in mRNA level in the wild type plants under B deficiency, and reduced mRNA accumulation for *BSU1*, no significant changes for *BES1* were observed in our RNA-seq result (Supplemental Fig. S2C-H). CONSTITUTIVE PHOTOMORPHOGENSIS AND DWARF (CPD) was up-regulated by B deficiency (Supplemental Fig. S2B). What’s more, the content of BL, the most active BR, was reduced under B deficiency in the wild type (En2) (Fig. 2D). Based on these results, we speculate that B deficiency is likely to negatively regulate BR signalling to affect root growth.

### BR signalling is involved in root responses to boron deficiency

BRs regulate root growth by regulating the elongation of differentiated cells and meristem size (Gonzalez-Garcia *et al*., 2011; Hacham *et al*., 2011; Fridman *et al*., 2014). To investigate the possible role of BRs in root responses to B deficiency, we grew BR signalling-related mutants under adequate (100 μM) and depleted (0.1 μM) B conditions and scored their primary root length. The *bes1-D* mutant, which over-accumulates the dephosphorylated active BR-regulated transcription factor BES1 (Yin *et al*., 2002), showed significantly longer primary root length than wild-type En2 under 0.1 μM B condition (Fig. 3A and B) and the difference was even more significant under 0.05 μM B condition (Fig. 3D; Supplemental Fig. S3A and B) or treated for longer time (17 DAG) under 0.1 μM B condition (Supplemental Fig. S3C and D). Quantification of root growth in the medium with B deficiency showed that *bes1-D* roots exhibited greater growth rate than wild-type under B deficiency (Fig. 3C). Cellular analysis revealed that *bes1-D* displayed longer mature cells, larger meristem and more meristematic cells than wild-type under B deficiency (Fig. 3E-G). To test whether BZR1 activity also impedes root response to low B stress, we established plant lines harbouring the same stabilizing mutation in BZR1 (*pBZR1-bzr1-D*). Like *bes1-D, pBZR1-bzr1-D* also showed significantly longer primary root length than wild-type under B deficiency condition (Supplemental Fig. S3E and F).

**Figure 3.**
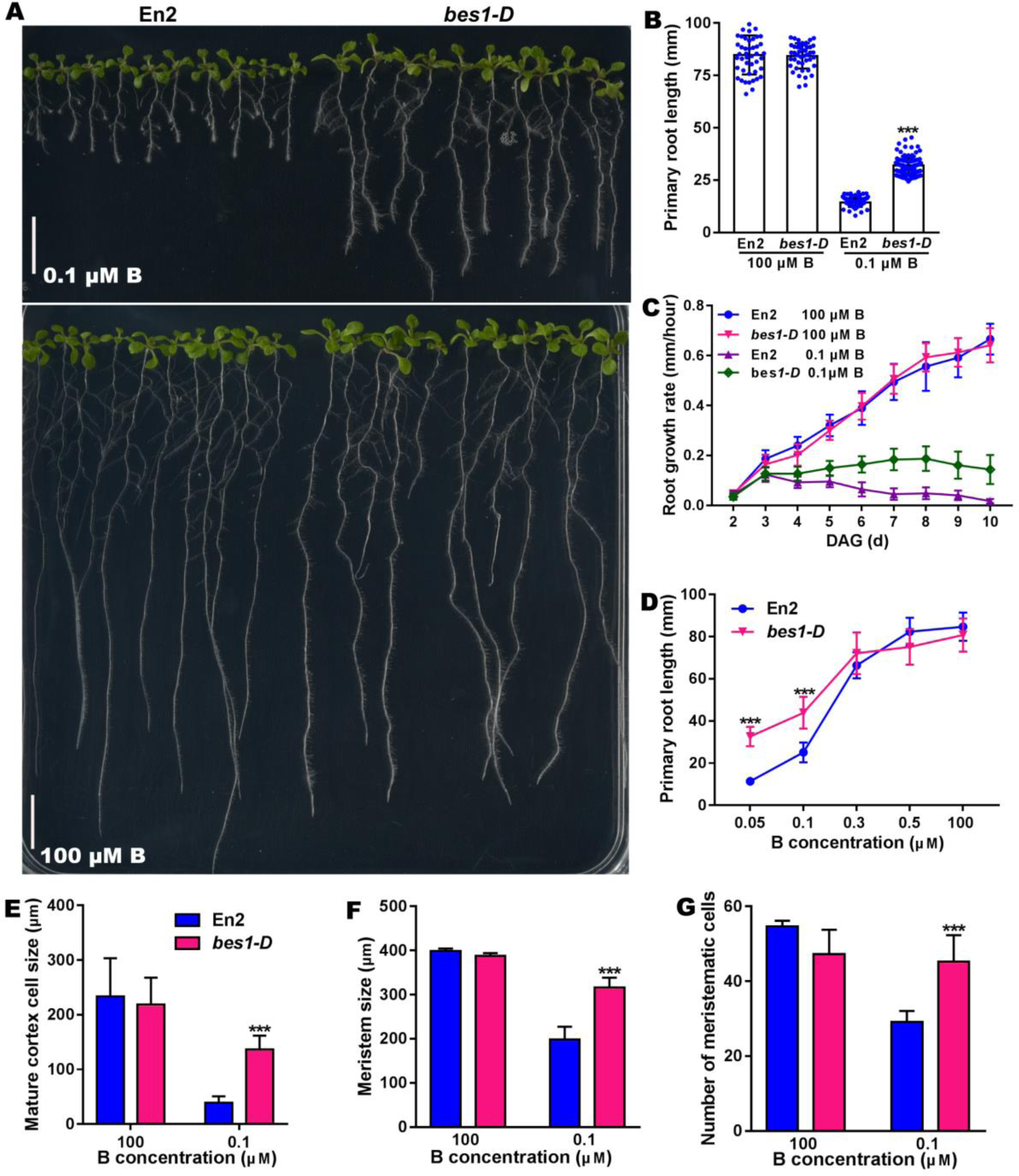
BES1 activity confers root insensitivity to B deficiency. **(A)** Phenotype of wild-type (En2) and *bes1-D* plants grown on MGRL media for 10 d under 0.1 and 100 μM B conditions. Scale bar, 10 mm. **(B)** Primary root length and **(C)** root growth rate of En2 and *bes1-D* plants grown as in **(A)** (mean ± s.d., n ≥ 45). The asterisks indicate a statistically significant difference (*t*-test, P < 0.001). **(D)** Root sensitivity of wild-type (En2) and *bes1-D* to increasing concentrations of B (mean ± s.d., n ≥45). The asterisks indicate a statistically significant difference (*t*-test, P < 0.001). **(E-G)** Mature cortex cell size **(E)** (mean ± s.d., n>40), meristem size **(F)** (mean ± s.d., n>6), number of meristematic cells **(G)** (mean ± s.d., n>6). The asterisks indicate a statistically significant difference (*t*-test, P < 0.001).

To strengthen the genetic evidence supporting the role of BR signalling in B deficiency mediated root growth, we used BR receptor mutant *bri1-301* (Xu *et al*., 2008), which showed shorter root than wild-type plants under B sufficient condition (Fabregas *et al*., 2013). However, the *bri1-301* mutant failed to respond to B depletion (Fig. 4A-D). These genetic evidences clearly suggest that BR signalling positively impacts on root responses to decreased B concentration.

**Figure 4.**
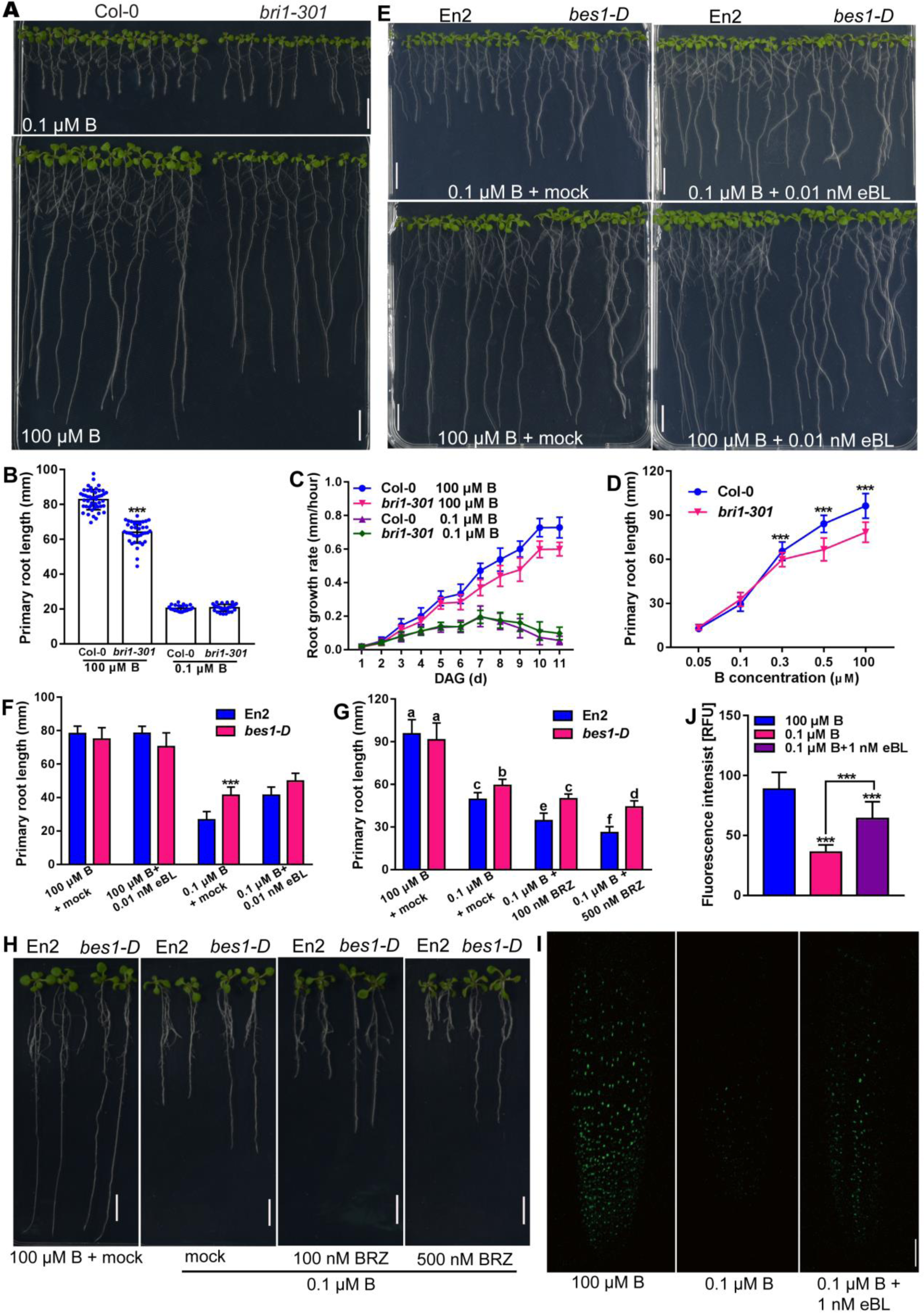
BR signaling is involved in root responses to boron deficiency. **(A)** Phenotype of wild-type (Col-0) and *bri1-301* plants grown on MGRL media for 9 days under 0.1 and 100 μM B conditions. Scale bar, 10 mm. **(B)** Primary root length of wild-type (Col-0) and *bri1-301* plants grown as in **(A)** (mean ± s.d., n ≥ 30). The asterisks indicate a statistically significant difference (*t*-test, P < 0.001). **(C)** Root growth rate of wild-type (Col-0) and *bri1-301* plants grown on MGRL media for 11 d under 0.1 and 100 μM B conditions (mean ± s.d., n ≥ 30). **(D)** Root sensitivity of wild-type (Col-0) and *bri1-301* to increasing concentrations of B (mean ± s.d., n ≥ 40). The asterisks indicate a statistically significant difference (*t*-test, P < 0.001). **(E)** Phenotype of wild-type (En2) and *bes1-D* plants grown on MGRL media for 9 d under 0.1 and 100 μM B conditions in the absence and presence of 0.01 nM eBL. Scale bar, 10 mm. **(F)** Primary root length of En2 and *bes1-D* seedlings grown as **(E)** (mean ± s.d., n ≥ 40). The asterisks indicate a statistically significant difference (*t*-test, P < 0.001). **(G)** Primary root length of En2 and *bes1-D* seedlings grown as **(H)** (mean ± s.d., n > 25). The different letters above the columns indicate significant differences between all genotypes and all growth conditions (Tukey-test, p ≤ 0.05). **(H)** Phenotype of wild-type (En2) and *bes1-D* plants grown on MGRL media for 10 days under 0.1 and 100 μM B conditions in the presence or absence of the BR biosynthesis inhibitor BRZ are shown. Scale bar, 10 mm. **(I)** A confocal image. Snapshots of the BES1-GFP signal, seedling transferred from adequate B medium to adequate B, B deficiency and B deficiency supplemented with eBL conditions for 24 h, respectively. Scale bar, 50μm. **(J)** Fluorescent signal measured in the seedling grown as **(I)** (mean ± s.d., n = 50). The asterisks indicate a statistically significant difference (*t*-test, P < 0.001).

BRs regulate plant root growth and development in a dose-dependent manner (Roddick *et al*., 1993; Clouse *et al*., 1996; Mussiget *et al*., 2003). If B deficiency downregulated BR signalling to inhibit root elongation, exogenous application of 24-epibrassinolide (eBL), a bioactive BR, should rescue the B deficiency-induced phenotype. Indeed, wild-type root growth was significantly enhanced in the presence of 0.01 nM eBL, which was similar to the length of *bes1-D* under B deficiency (Fig. 4E and F). While application of BR biosynthesis inhibitor BRZ aggravated root growth inhibition of wild-type under B deficiency condition, BRZ treatment did not affect *bes1-D* root’s insensitivity to B deficiency (Fig. 4G and H). BRs increase nuclear accumulation of both BES1 and BZR1 (Wang *et al*., 2002; Yin *et al*., 2002). In *p35s-BES1-GFP* plants, we found that nuclear signal was reduced significantly under B deficiency, while increased nuclear signal was observed following exogenous eBL application (Fig. 4I and J). Taken together, these results indicate that B deficiency downregulates BR signalling to inhibit root growth.

### BR signalling impacts neither B transport nor translocation

The root growth of *nip5;1* mutant was severely inhibited under B deficiency condition, and enhanced expression of *NIP5;1* and *BOR1* could improve the resistance to low B stress in *Arabidopsis* (Miwa *et al*., 2006; Kato *et al*., 2009). To assess whether BES1 activity can trigger *AtNIP5;1* gene expression, we analysed the relative expression of B transport or translocation genes in roots and shoots. In wild-type, the transcription level of *AtNIP5;1* in root was greatly higher under B deficiency condition than under B sufficient condition (Fig. 5A), consistent with previous studies (Takano *et al*., 2011). However, lower expression was observed in root of *bes1-D* plants under B deficiency condition (Fig. 5A). We also analysed other B-related genes, including *AtNIP6;1, AtNIP7;1, AtBOR1, AtBOR2*, and *AtBOR4*. Except *AtBOR1* had higher expression level in shoot of *bes1-D* mutant than wild-type under both B sufficient and deficient conditions, no obvious changes were observed in the transcription levels of *AtNIP6;1, AtNIP7;1, AtBOR2* and *AtBOR4* between the mutant and wild-type plants (Fig. 5B-F).

**Figure 5.**
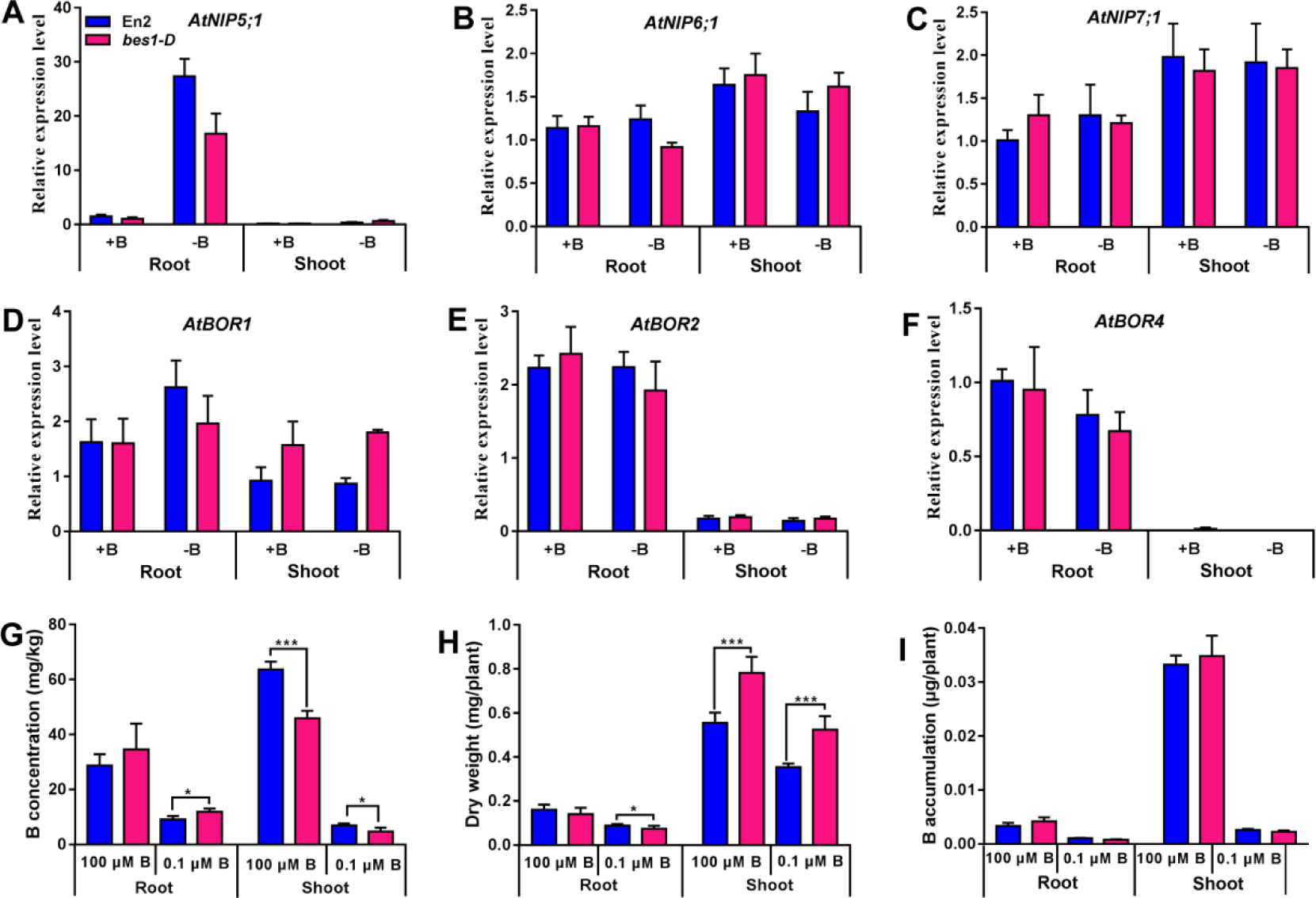
Relative expression of B transport-related genes, B homeostasis and dry weight in En2 and *bes1-D*. **(A-F)** Relative expressions of *NIP5;1* **(A)**, *NIP6;1***(B)**, *NIP7;1* **(C)**, *BOR1* **(D)**, *BOR2* **(E)**, *BOR4* **(F)** in root and shoot under 100 and 0.1 μM B (mean ± s.d., n=3). **(G-I)** B concentration, dry weight and B accumulation of En2 and *bes1-D* plants grown on MGRL media for 10 days under 0.1 and 100 μM B conditions (mean ± s.d., n=4). The asterisk shows a significant difference. *, P <0.05 with two-tailed Student’s t test. **, P <0.01 with two-tailed Student’s t test. ***, P <0.001 with two-tailed Student’s t test.

To indicate whether the expression changes of B-related genes are associated with the endogenous B homeostasis, we performed inductively coupled plasma-optical emission spectrophotometry (ICP-OES) analysis to quantify the B concentration and accumulation in both wild-type and *bes1-D* plants grown under adequate and depleted B conditions. We found *bes1-D* had higher B concentration than wild-type in root but lower in shoot (Fig. 5G), and it seemed to confer to the lower expression of *AtNIP5;1* in root of *bes1-D* plants under B deficiency condition (Takano *et al*., 2011). The difference in B concentration is likely to be caused by the different dry mass of *bes1-D* and wild-type (Fig. 5H). Furthermore, no significant differences in B accumulation were detected between wild-type and *bes1-D* mutant (Fig. 5I). Taken together, the insensitivity of *bes1-D* to B deficiency results from *BES1* acting independently of B transport or translocation pathway.

### Boron deficiency reduces the accumulation of BL

The mechanisms that B deficiency impinges BR signalling may be diverse. Based on the findings, we hypothesized that B deficiency reduces BR activity and/or levels to inhibit root growth. To address this hypothesis, we analyzed the relative expression levels of genes involved in BR biosynthesis in wild-type and *bes1-D* mutant under B sufficient and deficiency condition with qRT-PCR. Consisting with the result of RNA-seq, the expression levels of *CPD* genes were higher under B deficiency than sufficient B supply (Fig. 6A and Supplemental Fig. S2B). Interestingly, both *BR6ox1* and *BR6ox2* genes were down-regulated under B deficiency, in contrast with the negative feedback of reduced BR signalling (Fig. 6B and C; Supplemental Fig. S2I and J). *BR6ox2* involved in mediating the conversion of CS to BL in the rate-limiting final step of BRs biosynthesis (Kim *et al*., 2005). In *bes1-D* mutant, the expression levels of these genes were consistently lower than the wild type (Fig. 6A-C), which is consist with the known negative feedback regulation of *BES1* on their expression (Tanaka *et al*., 2005).

**Figure 6.**
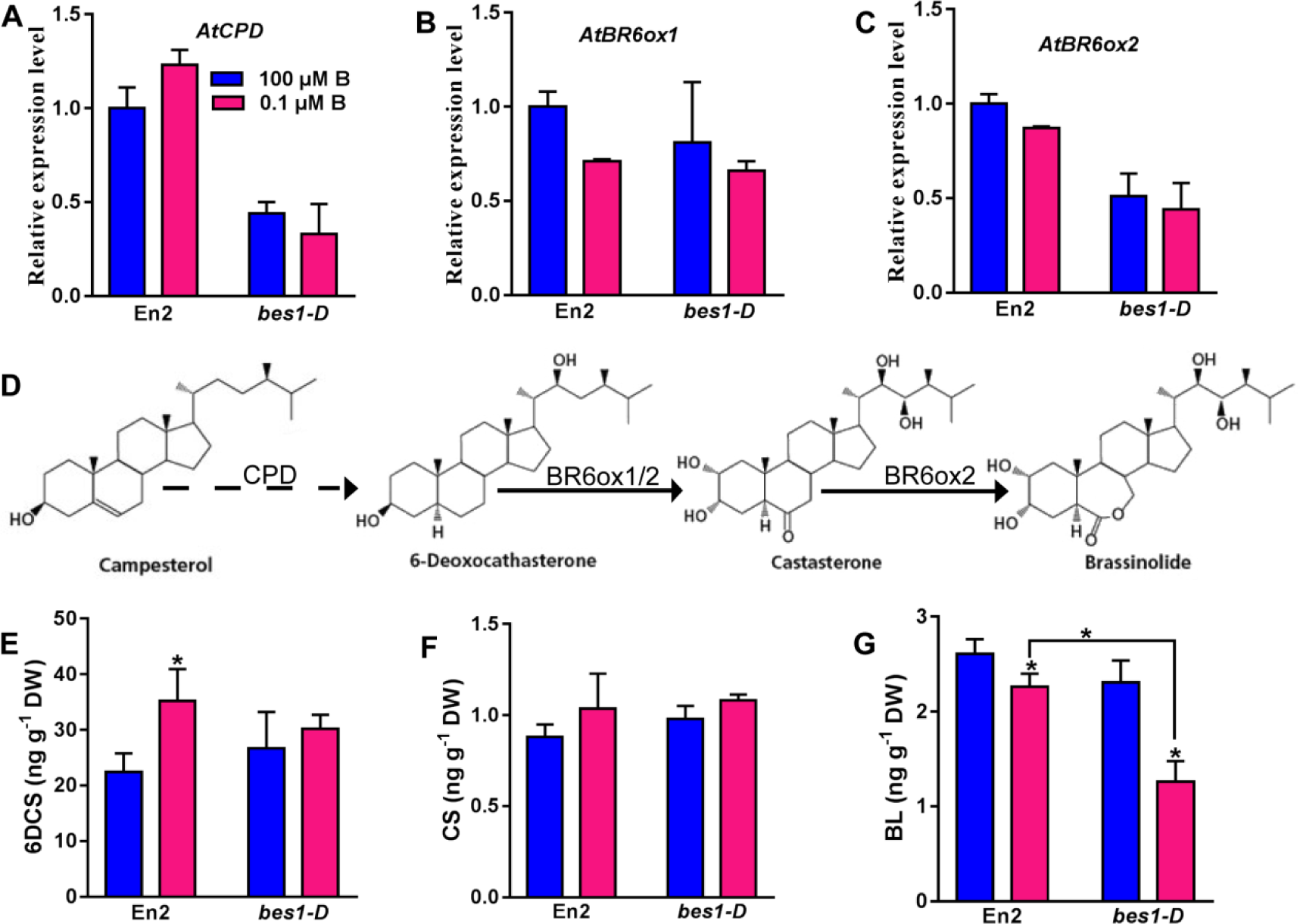
Low B stress triggers a reduction of BL. **(A-C)** Analysis of relative expression level of BR biosynthesis genes in roots of En2 and *bes1-D* plants grown 100 and 0.1 μM B (mean ± s.d., n=3). **(D)** Structures of BR molecules. **(E-G)** Content of detected 6DCS **(E)**, CS **(F)** and BL**(G)** are shown (mean ± s.d., n=3). The asterisk shows a significant difference. *, P <0.05 with two-tailed Student’s t test.

The significant changes in expression of BR biosynthesis genes prompted us to determine the endogenous levels of BRs in wild type and *bes1-D* plants. Several BR derivatives in both wild-type and *bes1-D* seedlings grown in B sufficient and deficiency conditions were analysed, including the most active BR, BL, and its immediate precursor, castasterone (CS), and the precursors of CS, 6-deoxocastasterone (6DCS). Indeed, BL levels were decreased in both wild-type and *bes1-D* plants under B deficiency condition (Fig. 6G and Supplemental Table S3), and *bes1-D* mutant accumulated less BL than wild-type, in agreement with that BL biosynthesis was under a negative feedback regulation of BES1 activity (Tanaka *et al*., 2005). No difference was found in CS level, but there were more 6DCS accumulated in wild-type under B deficiency (Fig. 6E and F; Supplemental Table S3). Both 6DCS and CS are synthesized from their corresponding higher order precursors (e.g. campesterol) by BR-specific synthetases like CPD, BR6ox1 and BR6ox2 (Miklós Szekeres *et al*., 1996; Shimada *et al*., 2003) (Fig. 6D). The accumulation of 6DCS was probably due to the up-regulation of *CPD* gene and down-regulation of *BR6ox1* and *BR6ox2* genes in wild-type under B deficiency, and the reduced *BR6ox2* gene expression conferred to the lower level of BL in wild-type under B deficiency. *BR6ox1* and *BR6ox2* should have been up-regulated by weakened BR signal through feedback pathway theoretically (Tanaka *et al*., 2005). It’s obvious that B deficiency disrupted this feedback, but the regulation network is unknown. Altogether, our data indicate that B deficiency impinges on BR signalling and BR-signalling dependent root growth at the transcriptional level of *BR6ox1* and *BR6ox2* genes.

## DISCUSSION

Although there’s been almost one century of research on its biology, B is still a much unknown micronutrient. B deficiency causes the inhibition of primary root growth rapidly (Fig. 1A-C), but mechanisms of root responses to B deficiency are poorly understood. Here, we used genetic, genomic, biochemical, and cell biological approaches to demonstrate that B deficiency impinges on BR signalling at the level of BL to control BR signalling-dependent root elongation.

The phytohormone BRs play pivotal roles in the regulation of plant growth and development with respect to environmental variability (Clouse & Sasse, 1998; Divi & Krishna, 2009). BRs regulate root growth by regulating the elongation of differentiated cells and meristem size with a dose-dependent manner (Chory *et al*., 1991; Gonzalez-Garcia *et al*., 2011; Fridman *et al*., 2014). B deficiency affects cell division and expansion, and finally results in root growth inhibition (Fig. 1D-I). We hypothesize that BR signalling may be involved in root responses to B deficiency. In agreement, first, 45.9% of the genes regulated by B deficiency in root were also regulated by BRs (Fig. 2B). We further analysed the expression patterns of these genes. Interestingly, more than 60% of the co-regulated genes response to B deficiency and BRs in the opposite directions (Fig. 2C and Supplemental Fig. S2A). Therefore, we speculate that B deficiency negatively regulates BR signalling to inhibit root growth. Second, we confirmed our hypothesis with different BR signalling mutants. Our data showed that both constitutive BR response mutant *bes1-D* and BR receptor mutant *bri1-301* were insensitive to B deficiency (Fig. 3A-D and Fig. 4A-D), and the enhanced root length of *bes1-D* mutant was the consequence of increased root cell expansion and proliferation (Fig. 3E-G). Third, exogenous application of eBL could rescue the root growth inhibition of wild-type to reach the length of *bes1-D* under B deficiency (Fig. 4E and F). While application of the BR biosynthesis inhibitor BRZ made wild-type plants more sensitive to B deficiency (Fig. 4G and H). Moreover, the nuclear signal of BES1-GFP was decreased under B deficiency, and could be restored by eBL treatment (Fig. 4I and J). In conclusion, our work provides complementary evidence demonstrating that B deficiency downregulates BR signalling to control the BR signalling-dependent root growth.

To further explore the effects of B deficiency on BR signal, we examined the homeostasis of endogenous BRs and the expression activity of BR synthetases genes. Under B deficiency, BL concentration in WT was decreased significantly (Fig. 6G), resulting in the up-regulation of *CPD* through negative feedback (Fig. 6A and Supplemental Fig. S2B). The up-regulation of *CPD* led to the increase of 6DCS (Fig. 6D and E). However, CS, the derivative of 6DCS, did not increase under B deficiency, which could be explained by the decrease of *BR6ox1* and *BR6ox2* expression (Fig. 6B-D, F and Supplemental Fig. S2I-J). The reduced expression of *BR6ox2* also contributed to the decline of BL concentration (Fig. 6C, D and G). In line with this supposition, BL concentration was even lower along with lower transcription activity of *BR6ox2* in *bes1-D*.

We speculated whether BR can trigger B-related genes expression to improve root elongation under B deficiency. The transcription level of *AtNIP5;1* in *bes1-D* is lower than wild-type in root (Fig. 5A), which is probably due to the higher concentration of B in *bes1-D* (Fig. 5G), because *NIP5;1* mRNA is accumulated under low B conditions but not under high B conditions, mediated by uORF of *NIP5;1* UTR region (Takano *et al*., 2006). There’s no significant difference in other B-related genes involved in B uptake and transport (Fig. 5B-F). Taken together, BES1 regulates primary root length under low B stress not by influencing the expression of B-related genes.

Previously, evidence about the involvement of auxin, ethylene, and cytokinin in the responses of root growth to B deficiency in *Arabidopsis* has been reported (Abreu *et al*., 2014; Juan *et al*., 2015; Poza-Viejo *et al*., 2018). Crosstalk between BR and other hormones has been observed at the physiological and transcriptional levels (Goda *et al*., 2004; Nemhauser *et al*., 2004; Hardtke *et al*., 2007; Gendron *et al*., 2008). Decreased BR signalling under B deficiency may have the ability to regulate other phytohormones signalling. Future work will elucidate whether reduced BR signalling under B deficiency interact with other phytohormones. We also noticed the primary root length of *bes1-D* was obviously longer than wild-type under B deficiency condition, but it reached only half the primary root length of wild-type under normal B condition (Fig. 3A). This may owe to the 54.1% genes regulated by B deficiency but not by BR (Fig. 2B). Future functional studies of the B deficiency regulated genes will further advance our understanding of the plant responses to B deficiency.

It is well known that the BL concentration in reproductive organs is significantly higher than that in vegetative organs, and the reproductive development of many crops, such as *Brassica napus*, is hypersensitive to B deficiency. Is the decrease of BL under B deficiency more considerable in reproductive organs? Does that suggest the potential role of BR signalling in B nutrition of plant reproduction? It needs further studies to answer these questions. Together, our work provides complementary evidence demonstrating that BRs involve in B deficiency-induced root reduction. Under B insufficient condition, the down-regulation of *BR6ox1* and *BR6ox2* leads to decreased BL biosynthesis, thus reduces BR signal and inhibits root elongation (Fig. 7). This is the framework of mechanism underlying BR’s involvement in root responses to B deficiency, but a deeper and more detailed understanding of the regulation pathway requires future studies.

**Figure 7.**
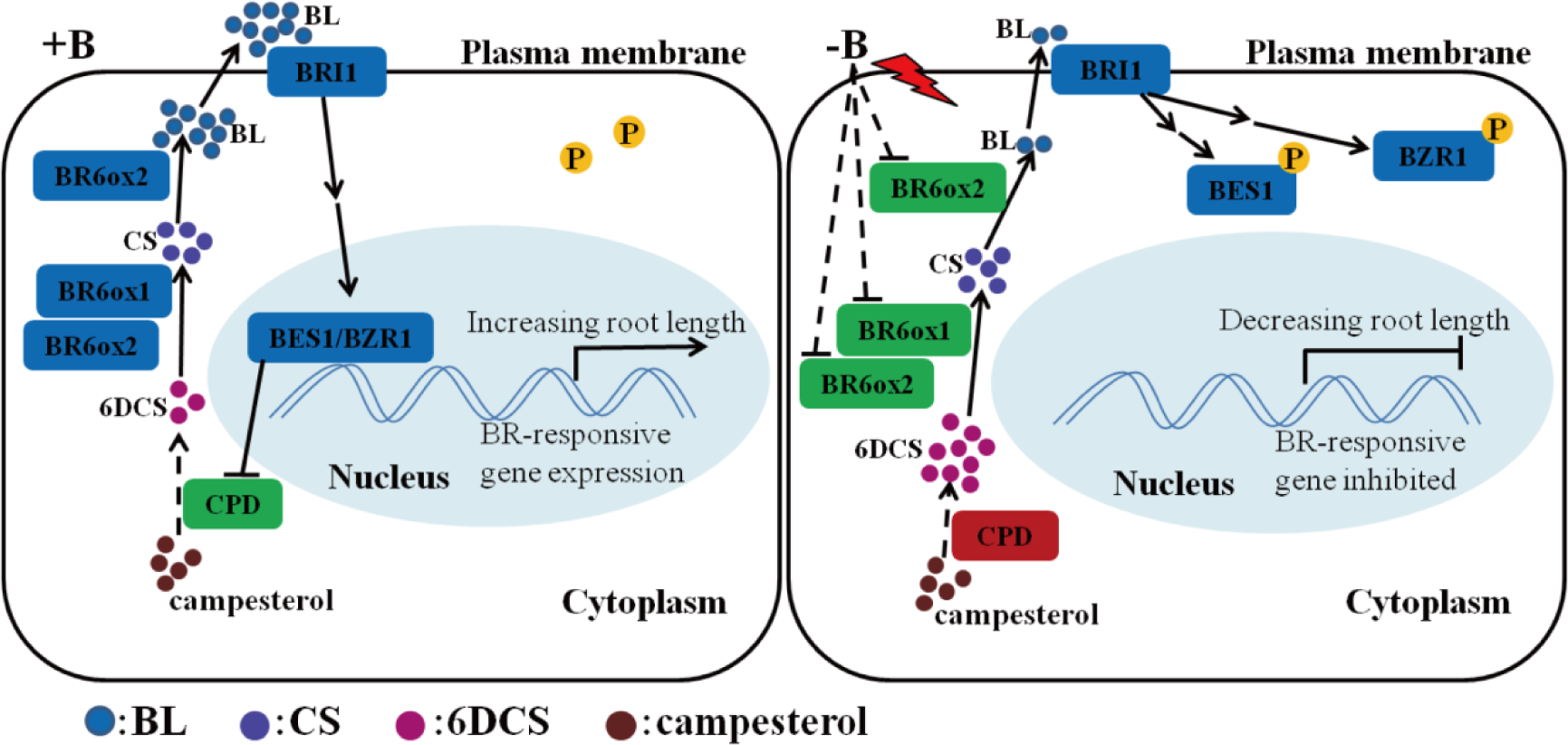
A model of boron deficiency-induced root growth inhibition is mediated by BRs in *Arabidopsis*. The BR biosynthesis genes, including *CPD*, is up-regulated by low B stress. *BR6ox1* and *BR6ox2*, are down-regulated by low B stress. The coloured red and green of rectangle indicate that gene are up-regulated and down-regulated under B deficiency, respectively. Under B sufficiency (left), the high concentration of BL result in nuclear accumulation of two unphosphorylated transcription factors, BZR1 and BES1, leading to BR-responsive gene expression and increasing root length. Compared with B sufficient conditions, B deficiency (right) reduced BL content by down-regulating the expression of *BR6ox1* and *BR6ox2*. BRI1 perceived low BL concentration lead to BR signaling decreased, BES1 and BZR1 were phosphorylated, and BR-responsive genes were inhibited, leading to decreasing root length.

## MATERIALS AND METHODS

### Plant material, growth conditions, and experiment treatments

The mutants and transgenic lines used in our study were previously reported, including *bes1-D* (Yin *et al*., 2002), *bri1-301* (Xu *et al*., 2008), *CYCB1;1:GUS* (Gonzalez-Garcia *et al*., 2011). *pBZR1-bzr1-D* and *p35S-BES1-GFP* transgenic plants were generated in *Arabidopsis* ecotypes En2 and Col-0 background through agrobacterium-mediated floral dipping method, respectively (Clough *et al*., 1998). Table S4 describes the primers used for constructing of these BR transgenic plants. Seeds were sterilized with 1% (w/v) NaClO for 10 min and rinsed completely in ultrapure water. Two days after vernalization at 4°C, the seeds were germinated on solid vertical MGRL medium (Fujiwara *et al*., 1992) containing 1% sucrose and 1% Gellan Gum supplemented with 100 μM or 0.1 μM B unless otherwise indicated. For chemical and hormone treatments, the seeds were planted on relevant supplemented MGRL medium and analyzed after 9 to 11 d, unless otherwise indicated. eBL and BRZ were dissolved in 100% dimethyl sulfoxide. BRZ was added at concentration of 100 nM and 500 nM. eBL was added at a concentration of 0.01 nM unless indicated otherwise. All plants grown at a temperature regime of 22/20°C (light/dark) under an 16/8 h (light/dark) photoperiod with a light intensity of 300–320 μmol·m^−2^·s^−1^.

### RNA extraction and expression analysis

Nine-day-old En2 and *bes1-D* mutant grown on MGRL solid medium with 100 μM B were transferred to 100 or 0.1 μM B medium for 24 h, and then harvested for RNA extraction with the RNA extraction kit (Promega, Madison, WI, United States), followed with reverse transcription using ReverTra Ace qPCR RT Master Mix with gDNA Remover kit (TOYOBO). Quantitative real-time PCR assays were performed under a Real-time PCR Detection System of QuantStudio™6 Flex (ABI, USA) in a 384-well plate format using SYBR Green PCR (TOYOBO, Osaka, Japan). *UBQ5* and *ACTIN* genes were used as two reference genes. Relative expression values were analyzed through the method of ΔΔ threshold cycle. Three biological replicates were performed for all reactions. The primers for qRT-PCR are listed in Supplemental Table S4.

### Root growth parameters

For the analysis of root growth rate, the position of root tips was recorded at the same time from 1 to 10 or 11-day-olds, and the root elongation in 24 h was measured. For sensitivity assays, the seeds were planted on MGRL medium containing different B concentrations as shown in the text. After 9 or 10 d, the primary root length was calculated on acquired images of seedlings by the ImageJ software. For mature and meristem zone cell size analysis, root tips of seedlings were imaged under a Leica TCS SP8 confocal laser scanning microscopes (www.leica-microsystems.com/home/). The roots were soaked with 10 μM FM4-64 about 3 min before observation. FM4-64 was viewed at an excitation wavelength of 488 nm, and the emission light was collected at 556 nm. Meristem size was defined as the distance from the quiescent center to the boundary of meristem where the cell of cortical is twice the size of the prior cell. Mature zone cell size was acquired via measuring the length of epidermal cells which showed the first root hair bulge. The cell divisions of root tips were observed with transgenic plants *CYCB1;1:GUS*. GUS histochemical staining was carried out using 5-bromo-4-chloro-3-indolyl-b-D-glucuronide as substrate (Gonzalez-Garcia *et al*., 2011). The ROS staining was performed according to the Jens Müller et al (Müller *et al*., 2015).

### RNA-Seq analysis

Six samples of mRNA (wild type En2 grown on MGRL medium containing 100 μM or 0.1 μM B, in three biological repetitions) were collected from roots of 9-day-old seedlings. The RNA-Seq experiments were carried out by Novogene. Library construction was implemented according to Illumina instructions and sequenced on a HiSeq 2000, to obtain 125 bp/150 bp paired-end reads. These reads were mapped to the *Arabidopsis thaliana* TAIR10 genome using Hisat2 v2.0.4. Transcript coordinates from the TAIR10 reference set were used to guide the alignment process. After alignment, HTSeq v0.9.1 was used to count the reads numbers mapped to each gene. The differential expression genes (DEGs) were analyzed by the DESeq R package (1.18.0). The Benjamini and Hochberg’s approach were used to adjust the resulting P-values for controlling the false discovery rate. Genes with an adjusted P-value <0.05 found by DESeq were defined as differentially expressed. The GOseq R package was used to implement Gene Ontology (GO) enrichment analysis of differentially expressed genes.

### Boron measurement

Seedlings of En2 and *bes1-D* were germinated in MGRL media for 11 d under 0.1 and 100 μM B conditions. Roots and shoots, harvested separately, were dried at 105°C for 30 min and then developed into constant weight at 65°C. Total dry weight and number of plants were measured for counting B concentration and accumulation. The samples were ground into powders with carnelian mortar. Adding 10 mL of 1 M HCl to shake in 250 rpm shaker for 2 hours, the B was lixiviated out. Lixiviate solution were filtered with filter paper and diluted 2 to 6 times. The B concentration was measured by inductively coupled plasma-optical emission spectrophotometry (Thermo Scientific).

### Confocal microscopy

The T_3_ homozygous seeds of *35S:BES1:GFP* were planted on MGRL medium for 4 days under 100 μM B conditions, and then transferred to adequate B (100 μM B), deficient B (0.1 μM B) and deficient B supplemented with 1 nM eBL medium for 24 h. Root tips of the seedlings were viewed on a Leica TCS SP8 confocal laser scanning microscopes (www.leica-microsystems.com/home/). GFP was viewed at an excitation and emission wavelength of 488 nm. For comparing the protein levels directly, the laser intensity and detection settings were controlled constantly. Fluorescence measurements in roots were performed using Leica LAS AF Lite software.

### BR analysis

For BR content, seedlings of En2 and *bes1-D* were germinated in MGRL media for 11 days under 0.1 and 100 μM B conditions, and whole plant were harvested. The determination of BR was performed by ProNetsBio. Samples of 200 mg freeze drying weight were grinded to powder in liquid nitrogen and extracted with ice-cold 80% methanol for 2 h and BL (OlChem), CS (OlChem), 6DCS (Sigma) as internal standards. After centrifugation, the samples were purified by Bond Elut preloaded columns and elution with methanol, and then further purified on strata-X columns and elution with methanol. Methanol solution was dried with nitrogen, dissolved in methanol, filtered through 0.22 μm membrane and then analyzed by HPLC-MS/MS.

### Statistical analysis

The values are means ± SD. Significant differences between treatments were analyzed based on two tailed unpaired Student’s *t*-tests, and *P*-values pf <0.05 were considered statistically significant. In multiple comparisons, Tukey-test was implemented at p ≤ 0.05.

## SUPPLEMENTAL DATA

**Supplemental Figure S1.** The ROS staining of wild-type En2 plants grown on MGRL media for 10 d under 100 and 0.1 μM B conditions.

**Supplemental Figure S2.** The BR-related genes response to boron deficiency in root in transcriptomes.

**Supplemental Figure S3.** BES1 and BZR1 activity confers root insensitivity to B deficiency.

**Supplemental Table S1.** Differential expression of *Arabidopsis* root genes at 0.1 μM and 100 μM H_3_BO_3_.

**Supplemental Table S2.** BR and B deficiency co-regulated genes in the root.

**Supplemental Table S3.** BR concentration in whole seedlings of wild-type En2 and mutant *bes1-D* grown in sufficient B (+B) and deficient B (-B) conditions for 11 d.

**Supplemental Table S4.** Primers used for cloning and qRT-PCR.

## ACKNOWLEDGEMENTS

We sincerely thank Xuelu Wang and Haijiao Wang (Huazhong Agricultural University) for kindly providing *bes1-D, bri1-301*, and wild type En2 seeds, and Guangjie Li (Chinese Academy of Sciences) for providing the seeds of *CYCB1;1∷GUS* transgenic plants.

